# Analysis of the hypothalamic oxytocin system and oxytocin receptor-expressing astrocytes in a mouse model of Prader-Willi syndrome

**DOI:** 10.1101/2022.08.15.503958

**Authors:** Ferdinand Althammer, Moritz Wimmer, Quirin Krabichler, Stephanie Küppers, Jonas Schimmer, Henning Fröhlich, Laura Dötsch, Matthew K. Kirchner, Javier E. Stern, Alexandre Charlet, Valery Grinevich, Christian P. Schaaf

**Affiliations:** Institute of Human Genetics, Heidelberg University, Heidelberg; Department of Neuropeptide Research in Psychiatry, Central Institute of Mental Health, Heidelberg University, Mannheim, Germany; Center for Neuroinflammation and Cardiometabolic Diseases, Georgia State University, Atlanta, GA, USA; Centre National de la Recherche Scientifique and University of Strasbourg, Institute of Cellular and Integrative Neuroscience, Strasbourg, France

## Abstract

Prader-Willi syndrome (PWS) is a neurodevelopmental disorder characterized by hyperphagia, obesity, developmental delay and intellectual disability. Studies suggest dysfunctional signaling of the neuropeptide oxytocin as one of the key mechanisms in PWS, and administration of oxytocin via intranasal or systemic routes yielded promising results in both humans and mouse models. However, a detailed assessment of the oxytocin system in mouse models of PWS such as the Magel2-deficient *Magel2*^*tm1*.*Stw*^ mouse, is lacking. In this study, we performed an automated counting of oxytocin cells in the entire paraventricular nucleus of the hypothalamus of *Magel2*^*tm1*.*Stw*^ and wild-type control mice and found a significant reduction in the caudal part, which represents the parvocellular subdivision. In addition, based on the recent discovery that some astrocytes express the oxytocin receptor (OTR), we performed detailed analysis of astrocyte numbers and morphology in various brain regions, and assessed expression levels of the astrocyte marker GFAP, which was significantly decreased in the hypothalamus, but not other brain regions in *Magel2*^*tm1*.*Stw*^ mice. Finally, we analyzed the number of OTR-expressing astrocytes in various brain regions and found a significant reduction in the nucleus accumbens of *Magel2*^*tm1*.*Stw*^ mice, as well as a sex-specific difference in the lateral septum. This study suggests a role for caudal PVN OT neurons as well as OTR-expressing astrocytes in a mouse model of PWS, provides novel information about sex-specific expression of astrocytic OTRs, and presents several new brain regions containing OTR-expressing astrocytes in the mouse brain.

## 1 Introduction

Prader-Willi syndrome (PWS) is characterized by infantile hypotonia, weight gain and overeating during childhood, as well as developmental delay and intellectual disability (1). Individuals with PWS lack the paternal copy of the protein-coding gene *MAGEL2*, albeit other relevant genes in the PWS gene locus such as *SNORD116* or *NECDIN* are typically also affected in PWS (2). Truncating point mutations of paternal *MAGEL2* cause a neurodevelopmental disorder that is molecularly and clinically related to PWS, i.e. Schaaf-Yang syndrome (3, 4). In order to study gene-related behavioral deficits, different *Magel2* knockout (KO) mouse lines have been generated. *Magel2* KO mice display unique behavioral patterns, including reduced levels of anxiety, lack of social discrimination (*Magel2*^*tm1Stw*^ mouse) (5), as well as altered exploration behavior and social interaction (*Magel2*^*tm1*.*1Mus*^ mouse) (6, 7), while hyperphagia seems to be less common (8, 9). The main difference between these two genetically engineered mice lines lies in the alteration of the *Magel2* gene sequence (2). In the *Magel2*^*tm1*.*1Mus*^ mouse, no transcript is produced due to deletion of the gene by homologous recombination. In the *Magel2*^*tm1*.*Stw*^ mouse, insertion of the LacZ β-gal gene results in expression of a fused, dysfunctional Magel2 protein. Thus, the *Magel2*^*tm1*.*Stw*^ and the *Magel2*^*tm1*.*1Mus*^ strains represent different transgenic models of PWS.

Already in the 1990’s, pioneering work by Dick Swaab revealed that pathological alterations in the hypothalamic oxytocin (OT) system could potentially underlie several of the observed symptoms of PWS including hyperphagia and weight gain (10). Swaab found that brains of individuals with PWS contained less OTergic neurons within the hypothalamus, especially in the caudal/posterior paraventricular nucleus (PVN) of the hypothalamus, where parvocellular OT (parvOT) neurons reside (11). In the years after Swaab’s initial discovery, a multitude of studies deciphering the precise role of OT in PWS followed using both animal models and human patients (6, 7, 12-23). One major study conducted in children with PWS highlighted that intranasal OT administration successfully restored suckling behavior (19) and a follow-up phase III trial of this experiment is currently underway NCT04283578. Although the hypothalamic OT system has been extensively studied using the *Magel2 KO* models, several key questions remain: i) Are there differences in the various *Magel2* KO mouse models with respect to the OT system? ii) Do *Magel2*-deficient mice display altered morphology of OT neurons (soma size, dendrite size or number of bifurcations)? iii) Do *Magel2*-deficient mice display altered OT receptor (OTR) levels and if so, what cell types (neurons, glia etc.) are affected? We recently demonstrated the existence of OTR-expressing astrocytes in the central amygdala (24) of mice and rats and hippocampal CA2 (25) of rats. However, a potential role of OTR-expressing astrocytes in *Magel2*-deficient mice has – to the best of our knowledge – not yet been considered.

In this study, we performed a thorough analysis of the hypothalamic and extra-hypothalamic OT system and analyzed morphology, immunofluorescent intensity and numbers of OT neurons in male and female WT and Magel2^*tm1Stw*^ mice. In addition, we performed semi-automated counting of OT cells within the entire PVN to obtain near absolute OT cell numbers for both groups. Next, we assessed OTR mRNA levels using RNAScope hybridization in various brain regions of interest. We also analyzed astrocyte number, morphology and intensity of the astrocytic marker GFAP. Finally, we performed RNAScope for OTR mRNA combined with immunohistochemical staining for the astrocyte marker glutamine synthetase (GS) and identified three novel brain regions harboring OTR-expressing astrocytes in the mouse brain, namely the nucleus accumbens (NAcc), the ventral tegmental area (VTA) and lateral septum (LS). We focused on these two regions given their important role of oxytocin-dependent modulation of social reward (26-28) and social behavior (29-32). We first analyzed OTR-expressing astrocytes in the central amygdala (CeA) in male and female WT and *Magel2*^*tm1*.*Stw*^ mice, and found no differences between the four groups. We next investigated the NAcc and LS, two brain regions, for which OTR-positive astrocytes have not been previously reported. Unexpectedly, we found OTR-expressing astrocytes in both brain regions, although in the LS, they were only present in female mice. Intriguingly, we observed a significant reduction of OTR-expressing astrocytes in the NAcc. Thus, this study highlights for the first time that a genetic condition (i.e. PWS or *Magel2* deletion) can affect astrocytic OTR levels and presents three previously unknown regions containing OTR-positive astrocytes, which can be used as potential targets in future behavioral studies. In addition, we describe sex-specific differences in oxytocin receptor-expressing astrocytes in the mouse brain, as shown by the lack of OTR-positive astrocytes in the LS of male mice.

## 2 Materials and methods

All performed experiments in mice were approved by Regierungspräsidium Karlsruhe (Baden-Württemberg) and carried out in agreement with the respective guidelines. At all time, animals had *ad libitum* access to food and water and all efforts were made to minimize suffering and the numbers of animals used for this study.

### Animals

We used C57BL/6 mice and *Magel2*^*tm1Stw*^ mice of both sexes (8 – 14 weeks old). Mice were housed in cages (2 per cage) under constant temperature (22 ± 2°C) and humidity (55 ± 5%) on a 12-h light cycle (lights on: 08:00-20:00).

### Immunohistochemistry

Mice were sacrificed using CO2 and were first perfused at a speed of 20mL/min with 0.01M PBS (200mL, 4°C) through the left ventricle followed by 4% paraformaldehyde (PFA, in 0.3M PBS, 200mL, 4°C), while the right atrium was opened with an incision. Brains were post-fixated for 24 hours in 4% PFA at 4°C and transferred into a 30% sucrose solution (in 0.01M PBS) at 4°C for 1-2 days. For immunohistochemistry, 50 µm slices were cut using a Leica Cryostat (CM3050 S) and brain slices were kept in 0.01M PBS at 4°C until used for staining. Brain sections were collected in series using the merging anterior commissural as an initial landmark so that consistent collecting of brain tissue across all animals could be achieved. Brain slices were blocked with 5% Normal Horse Serum in 0.01M PBS for 1h at room temperature. After a 15-min washing in 0.01M PBS, brain slices were incubated for 24hrs (48hrs in case of PS38) in 0.01M PBS, 0.1% Triton-X, 0.04% NaN_3_ containing different antibodies: 1:100 PS38 Neurophysin-I (gift from Harold Gainer), 1:1000 anti-glutamine synthetase (monoclonal mouse, abcam, 176562), anti-GFAP (goat polyclonal, abcam, ab53554) at room temperature. Following 15-min washing in 0.01M PBS, sections were incubated in 0.01M PBS, 0.1% Triton-X, 0.04% NaN_3_ with 1:500 Alexa Fluor 488/594-conjugated donkey anti-rabbit/goat/mouse (Jackson ImmunoResearch, 711-585-152, 705-585-147, 715-545-151) for 4 hours at RT. Brain slices were washed again for 15 mins in 0.01M PBS and mounted using antifade mounting medium (Vectashield with DAPI, H-1200B).

### RNAScope

RNAScope reagents were purchased from acdbio (Multiplex v2 RNAScope kit). Nuclease-free water and PBS were purchased from Fisher Scientific. Brains were processed as described under ***Immunohistochemistry*** using nuclease-free PBS, water, PBS and sucrose. We followed the manufacturer protocol with a few modifications: 1) Immediately after cryosectioning, slices were washed in nuclease-free PBS to remove sucrose and OCT compound. 2) Hydrogen peroxide treatment was performed with free floating sections prior to slice mounting. 3) Sections were mounted in nuclease-free PBS at room temperature. 4) Pretreatment with Protease III was performed for 20 minutes at room temperature. 5) No target retrieval step was performed.

### Confocal microscopy and 3D IMARIS analysis

Confocal images were obtained using a Nikon C2 confocal microscope (1024×1024 pixel, 16-bit depth, pixel size 0.63-micron, zoom factor 1). For the three-dimensional reconstruction 50µm-thick sections (z-stacks) were acquired using 1µm-steps. Three-dimensional reconstruction of astrocyte somata and subsequent analysis of surface and filament reconstructions was performed as previously described (33). Image processing, three-dimensional reconstruction and data analysis were performed in a blind manner in regards to the experimental conditions.

### Analysis of OTR mRNA levels

Density and intensity of OTR immunosignal were assessed in Fiji (NIH) using the threshold paradigm. Average pixel intensity was calculated using the Analyze<Measure function using collapsed 50µm sections (z-stack) images. For density analyses, raw images were opened and threshold was set individually for each image to achieve a near-perfect overlap of raw immunosignal and pixels used for quantifications (**Figure S1**). Optical density (area fraction in %) was then calculated via the Analyze<Measure function using the thresholded images so that only pixels above the threshold are included in the further image analysis. For quantifications we used at least 4-6 sections for each animal and brain region, if not indicated otherwise.

### Statistical analysis

All statistical analyses were performed using GraphPad Prism 9 (GraphPad Software, California, USA). The significance of differences was determined using two-tailed Student’s t test, one-way or two-way ANOVA or one-sample t-test, as indicated in the respective figure legends. Chi-Square tests were used to compare differences in the incidence of OTR+ cells between sham and HF rats. Results are expressed as mean ± standard error of the mean (SEM). Results were considered statistically significant if p<0.05 and are presented as * for p<0.05, ** for p<0.01 and *** for p<0.0001 in the respective Figures.

## 3 Results

The hypothalamic OT system is mainly composed of two distinct nuclear formations, namely the supraoptic (SON) and the paraventricular nucleus (PVN) of the hypothalamus, albeit the anterior commissural, periventricular nucleus, the different accessory nuclei and other structures harboring OT neurons make up between 20-35% of all OT cells both in rats (11) and mice (34). As a brain-wide analysis of OT neuron numbers and morphology and OTergic innervation in various forebrain regions has not yet been performed for the *Magel2*^*tm1*.*Stw*^ mouse, we aimed to fill this gap. A reliable, automated analysis of SON OT neurons using the Imaris software was not feasible, given that OT neurons within the SON tend to cluster too densely. However, conventional analysis using 8-10 SON snapshots and manual counting of SON neurons, as well as analysis of their morphology (i.e. cell volume of somata) revealed no apparent differences between WT and *Magel2*^*tm1*.*Stw*^ mice (referred to as KO on the Figures) in males or females (data not shown). Similarly, analysis of cell numbers and morphology of OTergic neurons within the accessory nuclei revealed no apparent differences between the two groups (data not shown). Thus, we next focused on the PVN and analyzed the entire nucleus in WT and *Magel2*^*tm1*.*Stw*^ mice (**Figure 1A, B**, n=4 per group, both sexes) and assessed cell numbers (**Figure 1C, D**, divided into rostral, medial and caudal parts), immunofluorescent intensity and morphology (total cell volume in µm^3^). For the automated cell counting (**Figure 1F**), we did not differentiate between PVN and periventricular nucleus, which might explain why our numbers are slightly higher than what has been recently reported elsewhere (34). While we observed no difference in OT immunoreactivity (**Figure 1E**), both male and female *Magel2*^*tm1*.*Stw*^ mice displayed lower total OT cell numbers (**Figure 1G**). Automated cell counting via the Imaris software revealed significantly lower OT cell numbers for the caudal, but not the medial or rostral PVN (**Figure 1G**). Interestingly, this is in stark contrast to what has been previously reported for the *Magel2*^*tm1*.*1Mus*^ mouse, suggesting that the different genetic alterations may differentially impact OT neuron ontogenesis (35, 36). Already in the 1990’s Swaab reported a reduction of OT-positive cells in the brains of PWS patients (10), particularly in the posterior PVN, where parvocellular OT (parvOT) neurons reside in the mouse and rat brain (11). Thus, we speculated that the reduction in the total number of OT cells could be due to the specific absence of parvOT neurons within the caudal PVN and three-dimensionally reconstructed the surface of OT neurons to analyze cell volume of individual cells. Analysis of cell volume revealed no significant differences between WT and *Magel2*^*tm1*.*Stw*^ mice (**Figure 1H**), potentially arguing against this hypothesis. However, given that the vast majority of OT cells within the caudal part of the PVN (parvocellular subdivision) is parvocellular in mice (37), it seems plausible that a reduction of OT cells within this particular subdivision would not be reflected in our data set containing the cell volume of all OT cells. Thus, a more appropriate approach would be the intraperitoneal injection of the retrograde tracer Fluorogold™, which labels parvocellular, but not magnocellular OT neurons (11, 37), in order to quantify the absolute number of parvocellular OT cells in WT and *Magel2*^*tm1*.*Stw*^ mice.

**Figure 1.**
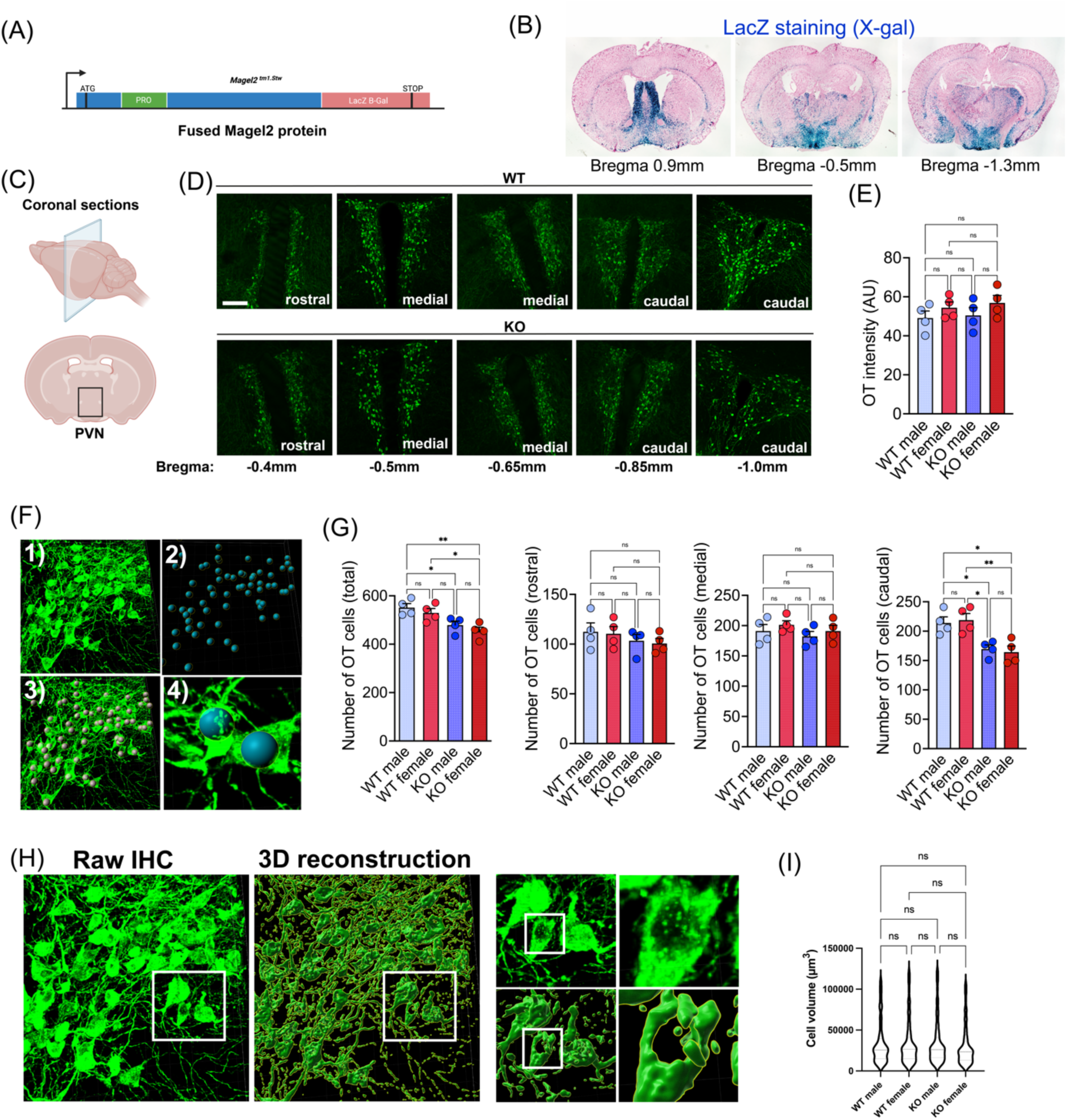
**(A)** Schematic depiction of the transgenic modification in *Magel2*^*tm1*.*Stw*^ mice; insertion of the LacZ B-Gal gene results in expression of a fused *Magel2* protein **(B)** Images show presence of LacZ reporter (X-gal staining) in *Magel2*^*tm1*.*Stw*^ mice. **(C)** Schematic depiction of coronal sectioning and anatomical location of the PVN. **(D)** Side-to-side comparison of OT neurons in the PVN of WT and *Magel2*^*tm1*.*Stw*^ mice at various Bregma levels. Scale bar=100µm. **(E)** No difference in OT signal intensity between WT and *Magel2*^*tm1*.*Stw*^ mice (n=5 per group). **(F)** Imaris-assisted, automated cell counting of PVN OT neurons via the spot function. (1) raw image of PVN OT neurons, (2) each spot represents one OT neuron, (3) co-localization of OT neurons and spots, (4) high magnification image showing placement of spots in the OTergic soma. **(G)** Quantification of OT cell numbers within the PVN in male and female mice. Significant differences were observed in the total number of OT cells and the caudal PVN (parvocellular subdivision, n=5 per group). **(H)** Depiction of the pipeline for three-dimensional reconstruction of OT neurons based on fluorescent immunolabeling. High magnification insets highlight the detailed reconstruction of OTergic somata. **(I)** No difference in OTergic morphology between WT and *Magel2*^*tm1*.*Stw*^ mice (n=5 per group). *p<0.05, **p<0.01, one-way ANOVA.

We were next interested to see whether global levels of OTRs varied between WT and *Magel2*^*tm1*.*Stw*^ mice and performed RNAScope hybridization for OTR mRNA using brain sections containing brain regions (NAcc, LS, central amygdala (CeA), insular cortex (InC) etc.), in which prominent OTR expression has previously been reported (38, 39). We first imaged the entire section using a slide scanner with 20x magnification (dry objective) to be able to verify that our signal matches with previously published data (**Figure 2A-D**). We observed clearly confined OTR mRNA signals in brain regions know to express OTRs in mice such as the NAcc (**Figure 2A**), LS (**Figure 2B**), IC (**Figure 2C**) or CeA (**Figure 2D**). Next, we assessed OTR mRNA levels (confocal imaging with 40x oil objective, area fraction in %) via thresholding analysis (**Figure S1**) in WT and *Magel2*^*tm1*.*Stw*^ mice (n=5 per group and sex) in 8+ brain regions. We only found significant changes in the NAcc and CeA, while the remaining brain regions appeared unaffected by the *Magel2* deletion (not shown). In the NAcc, *Magel2*^*tm1*.*Stw*^ mice displayed significantly lower OTR mRNA levels in both males and females (**Figure 2E**), while we observed the exact opposite in the CeA of male and female *Magel2*^*tm1*.*Stw*^ mice (**Figure 2F**).

**Figure 2.**
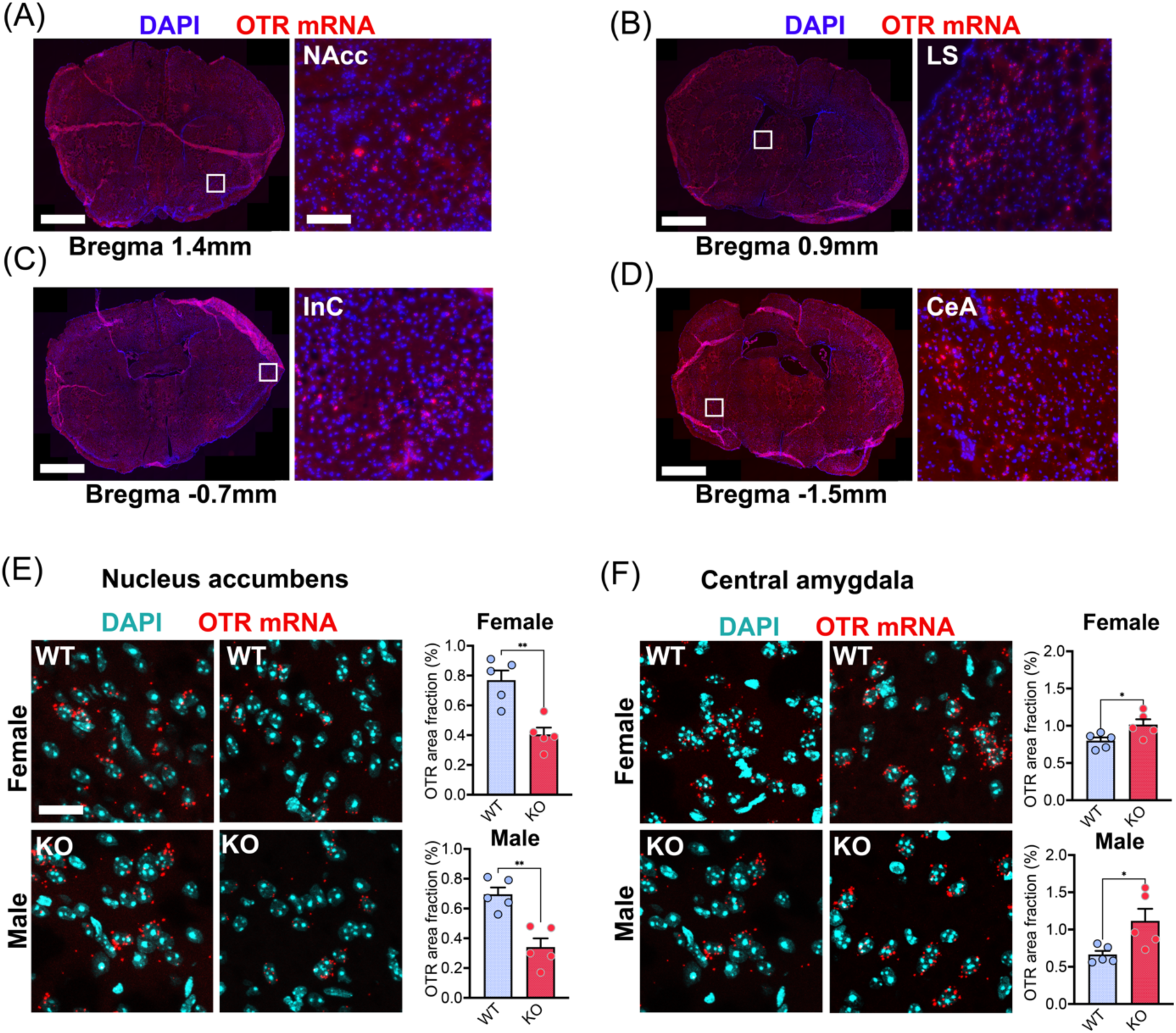
**(A-D)** Slide scanner images at 20x magnification showing whole brain sections of a WT mouse at various Bregma levels. High magnification insets depict NAcc **(A)**, LS **(B)**, InC **(C)** and CeA **(D)**. Size of brain sections was adjusted for aesthetic reasons, scale bars = 1.8mm **(A)**, 100µm (inset **(A)**, same for all others), 2.0mm **(B)**, 2.3mm **(C)** and 2.4mm **(D). (E)** Confocal images show representative examples of OTR mRNA levels in the NAcc of female and male WT and *Magel2*^*tm1*.*Stw*^ mice. OTR mRNA levels are significantly reduced in *Magel2*^*tm1*.*Stw*^ mice of both sexes compared to their WT controls (n=5 per group). **(F)** Confocal images show representative examples of OTR mRNA levels in the CeA of female and male WT and *Magel2*^*tm1*.*Stw*^ mice. OTR mRNA levels are significantly reduced in *Magel2*^*tm1*.*Stw*^ mice of both sexes compared to their WT controls (n=5 per group). Scale bar=25µm. *p<0.05, **p<0.01, one-way ANOVA.

The role of astrocytes in Prader-Willi syndrome is not well understood and only a handful of studies addressed this topic (40). Thus, we were interested to see whether *Magel2*^*tm1*.*Stw*^ mice display altered astrocyte number and/or morphological changes. To this end, we stained brain sections of WT and *Magel2*^*tm1*.*Stw*^ mice containing CeA, NAcc, LS and VTA against the astrocyte-specific marker glutamine synthetase (GS, **Figure 3A**). Given that GS predominantly stains the astrocytic soma (whereas GFAP only stains major astrocytic processes), it is a great tool to study astrocytic soma size. We did not observe significant changes in astrocytic cell numbers (i.e. GS-positive cells) in any of the four assessed brain regions across the four different groups suggesting that there is no permanent pathological proliferation or reduction in *Magel2*^*tm1*.*Stw*^ mice (**Figure 3B**). In addition, we performed three-dimensional reconstruction of astrocyte somata in all four brain regions across all four groups, but did not observe any significant changes (**Figure 3C, D**). Thus, we concluded that neither the number nor the soma size of astrocytes is changed in *Magel2*^*tm1*.*Stw*^ mice.

**Figure 3.**
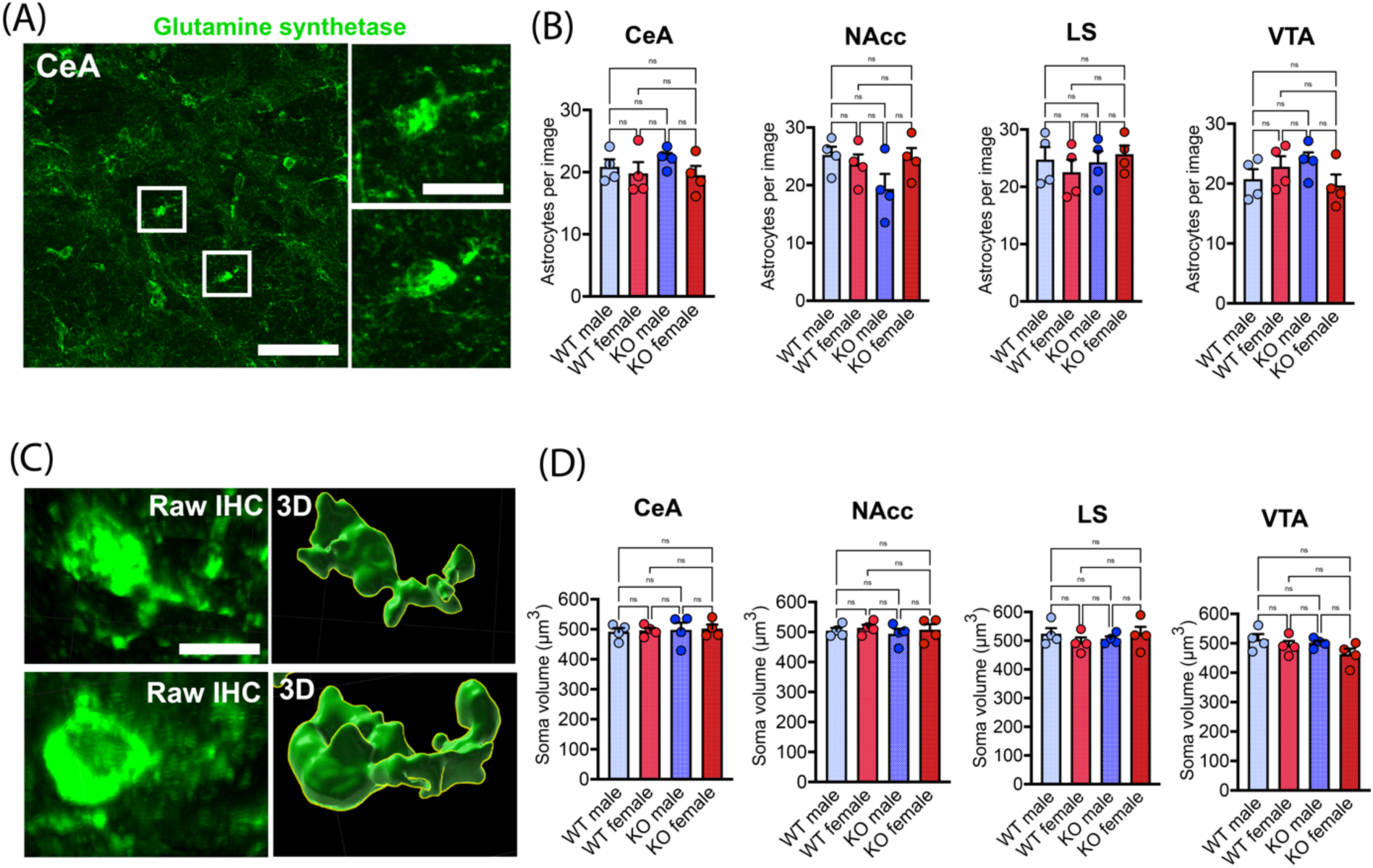
**(A)** Confocal image of GS-positive astrocytes in the CeA of a WT female. Scale bar=100µm and 20µm (enlarged inset). **(B)** Quantification of astrocyte numbers in the CeA, NAcc, LS and VTA in male and female WT and *Magel2*^*tm1*.*Stw*^ mice (n=4 per group). **(C)** Three-dimensional reconstruction of astrocyte somata via Imaris. **(D)** Quantification of astrocyte soma size (somatic volume in µm^3^) in the CeA, NAcc, LS and VTA in male and female WT and *Magel2*^*tm1*.*Stw*^ mice (n=4 per group).

Several studies suggest that genetic alterations within the PWS locus 15q11.2-q13 could promote low-grade systemic inflammation, as increased brain age (41) and elevated levels of cytokines (42, 43) have been reported in PWS patients. Since GFAP upregulation and astrocytic hypertrophy is one of the hallmarks of neuroinflammation in the brain (44-46), we sought to assess GFAP intensity in WT and *Magel2*^*tm1*.*Stw*^ mice. Given that GFAP expression is extremely variable and depends on various factors including age, brain regions, sex and experimental conditions (47-50), we tried to overcome this issue by using age-matched littermates for immunohistochemical analysis and by drastically increasing the number of images taken per animal. To this end, we obtained 4-6 image blocks (4*4=16 taken at 40x magnification) that were fused to one large image during post-processing, which amounted to 64-96 individual images per animal and brain region used for image analysis, respectively. We chose the hypothalamus, CeA and NAcc, as all these brain regions display substantial *Magel2* expression (51) and are involved in the modulation of various aspects of social behavior (26, 52, 53). To our surprise, analyzing these brain regions, we found that GFAP immunoreactivity in the hypothalamus (**Figure 4A-C**), but not the CeA (**Figure 4D-F**) or NAcc (**Figure 4G-I**) was significantly decreased. Interestingly, these findings are in line with a recent study (54), in which the authors describe decreased GFAP levels in the hypothalamus of PWS patients. Given that GFAP only stains a fraction (15-20%) of the astrocytic cytoskeleton and thus might not be the best tool for a detailed assessment of astrocytic morphology (55), we did not attempt to three-dimensionally reconstruct GFAP-labeled astrocytes in WT or *Magel2*^*tm1*.*Stw*^ mice.

**Figure 4.**
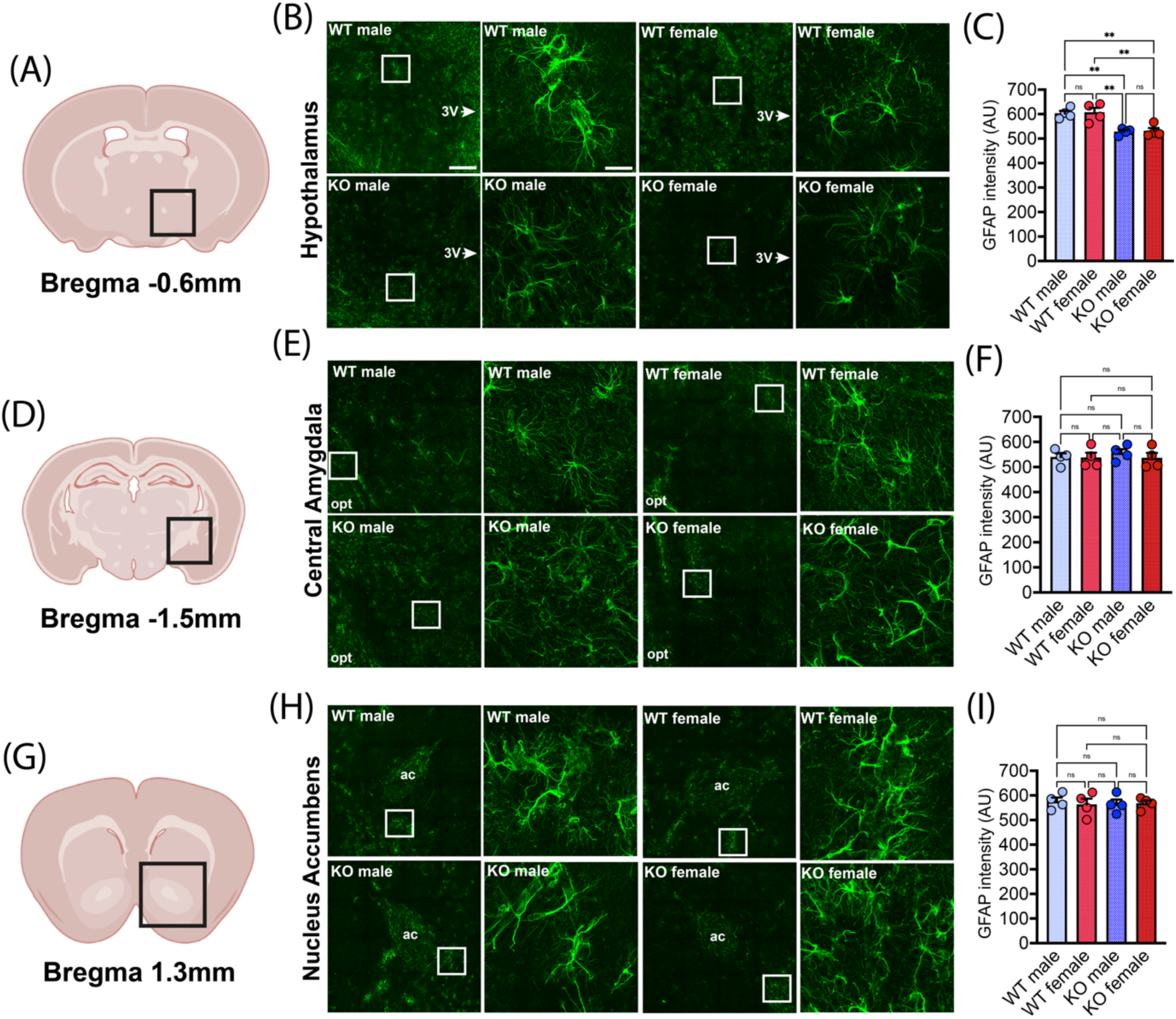
**(A)** Schematic depiction of the anatomical location of medial and ventral hypothalamus used for the analysis. **(B)** Representative confocal images of female and male WT and *Magel2*^*tm1*.*Stw*^ mice (whole hypothalamus, 16×40x confocal image, stitched). Enlarged insets show GFAP-positive astrocytes. 3V=third ventricle. **(C)** Significant reduction in GFAP intensity in the hypothalamus of male and female *Magel2*^*tm1*.*Stw*^ mice (n=4 per group). *p<0.05, **p<0.01, one-way ANOVA. **(D)** Schematic depiction of the anatomical location of the CeA. **(E)** Representative confocal images of female and male WT and *Magel2*^*tm1*.*Stw*^ mice (whole hypothalamus, 16×40x confocal image, stitched). Enlarged insets show GFAP-positive astrocytes. 3V=third ventricle. **(F)** No difference in GFAP intensity in the CeA between male and female WT or *Magel2*^*tm1*.*Stw*^ mice (n=4 per group). **(G)** Schematic depiction of the anatomical location of the NAcc. **(H)** Representative confocal images of female and male WT and *Magel2*^*tm1*.*Stw*^ mice (whole hypothalamus, 16×40x confocal image, stitched). Enlarged insets show GFAP-positive astrocytes. 3V=third ventricle. **(I)** No difference in GFAP intensity in the NAcc between male and female WT or *Magel2*^*tm1*.*Stw*^ mice (n=4 per group).

The presence and function of OTR-expressing astrocytes in the CeA has been described (24). In addition, we recently reported OTR-positive astrocytes in dorsal CA2 in rats (25), which seemed to be affected by myocardial infarction. Thus, we were curious to investigate potential alterations to OTR-positive astrocytes in our transgenic *Magel2*^*tm1*.*Stw*^ mouse model of PWS. We chose the CeA, NAcc and LS as brain regions of interest due to their important roles in exploratory behavior and anxiety (CeA), social reward (NAcc) and social behavior (LS). Given that *Magel2*^*tm1*.*Stw*^ displayed abnormal behavioral phenotypes that are potentially related to these brain regions – some of which could be rescued by OT administration – we hypothesized that the numbers of OTR-positive astrocytes within these brain regions might be altered in *Magel2*^*tm1*.*Stw*^ mice. For the CeA, we processed brain sections of male and female mice (n=3 per group and sex) via RNAScope for OTR mRNA with combined immunohistochemical staining against astrocyte markers GFAP and GS and analyzed the percentage of OTR-positive astrocytes (**Figure 5A-C, Figure S2A**). We observed that 12-25% of astrocytes were OTR-positive across all conditions, which is consistent with our previous results (24). We did not find any statistically significant differences between WT or *Magel2*^*tm1*.*Stw*^ mice (**Figure 5D**). We next analyzed brain sections containing the NAcc and – to our big surprise – observed OTR-expressing astrocytes in male and female brains of WT mice (**Figure 6A-C, Figure S2B**). Even more intriguingly, we found that the number of OTR-positive astrocytes was significantly reduced in brains of male and female *Magel2*^*tm1*.*Stw*^ mice (**Figure 6D**). Finally, we analyzed brain sections containing the LS of male and female mice (**Figure 7A-C, Figure S2C**), a region where – to the best of our knowledge – no OTR-expressing astrocytes have been reported yet in mice. Intriguingly, we found OTR-positive astrocytes only in the LS of female WT and *Magel2*^*tm1*.*Stw*^ mice (**Figure 7B, C**), but not their male counterparts, indicating a potential sex-specific difference in astrocytic OTR expression in this brain region (**Figure 7D**). In addition to LS and NAcc, we also found OTR-expressing astrocytes in the VTA, but poor tissue quality and fragility of very caudal sections did not allow us to perform a thorough analysis in WT and *Magel2*^*tm1*.*Stw*^ mice (**Figure S3A-C**). However, this is – to the best of our knowledge – the first account of OTR-positive astrocytes in the VTA of male and female mice.

**Figure 5.**
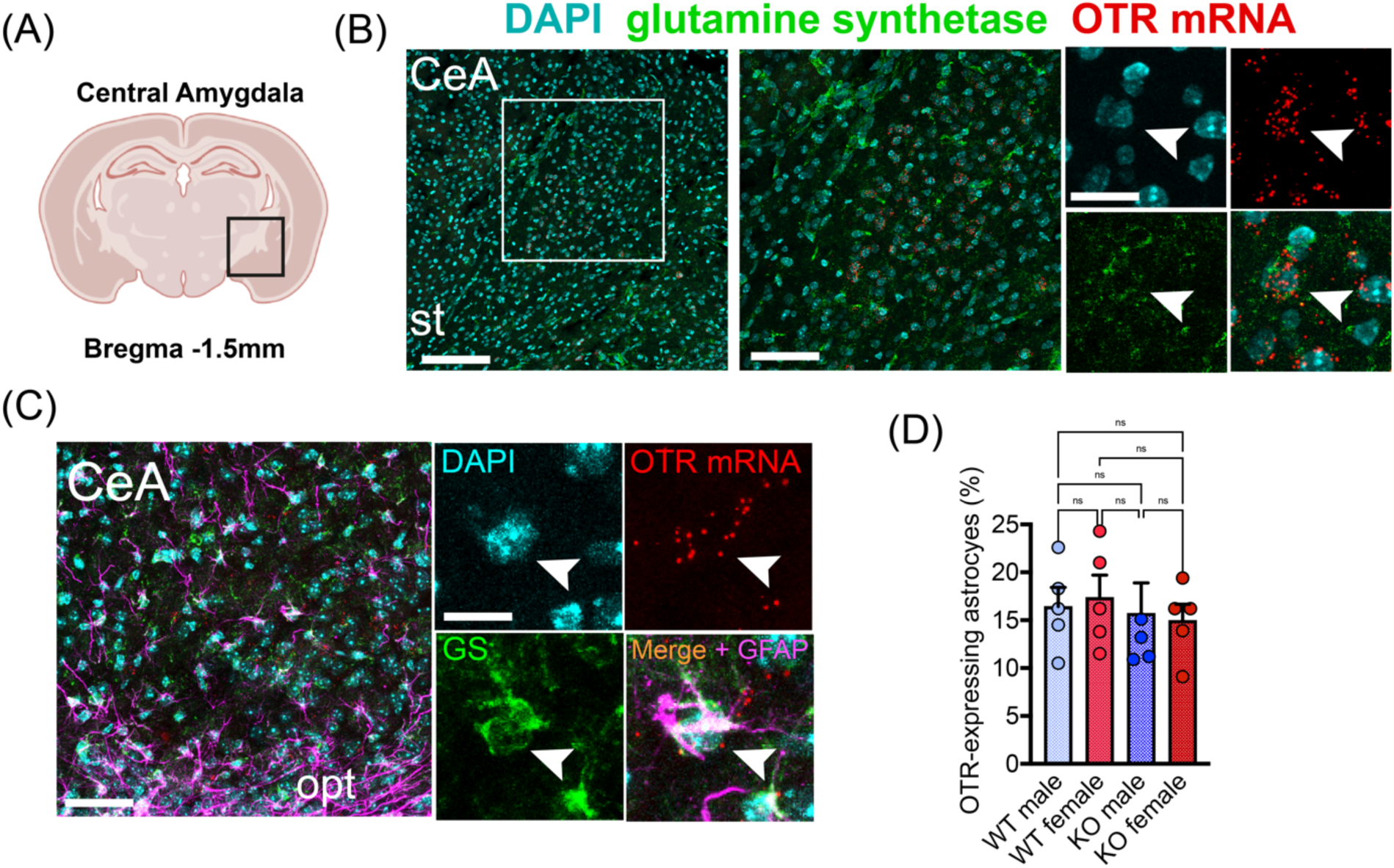
**(A)** Anatomical location of the CeA. **(B)** Confocal images showing RNAScope hybridization for OTR mRNA combined with immunostaining against the astrocyte marker GS. Enlarged inset shows colocalization of OTRmRNA and GS signals. Scale bar=200µm, 100µm and 25µm, st=stria terminalis. **(C)** Confocal images show RNAScope hybridization for OTR mRNA combined with immunostaining against the astrocyte markers GS and GFAP. Scale bar=100µm and 10µm. **(D)** No difference in the number of OTR-positive astrocytes between male and female WT and *Magel2*^*tm1*.*Stw*^ mice (n=5 per group).

**Figure 6.**
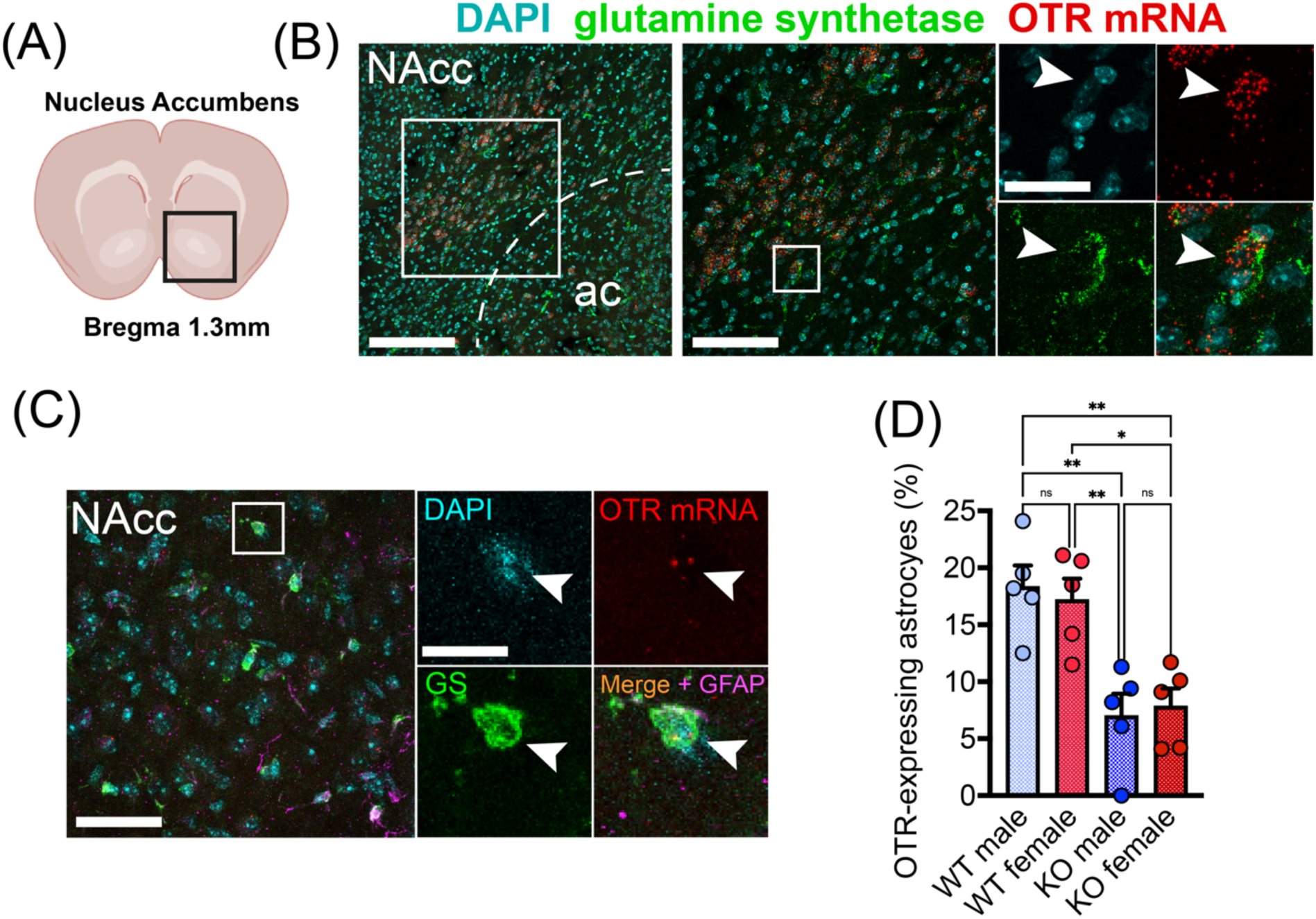
**(A)** Anatomical location of the NAcc. **(B)** Confocal images showing RNAScope hybridization for OTR mRNA combined with immunostaining against the astrocyte marker GS. Enlarged inset shows co-localization of OTRmRNA and GS signals. Scale bar=200µm, 100µm and 20µm, ac=anterior commisura. **(C)** Confocal images show RNAScope hybridization for OTR mRNA combined with immunostaining against the astrocyte markers GS and GFAP. Scale bar=100µm and 10µm. **(D)** Significant reduction in the number of OTR-positive astrocytes in male and female *Magel2*^*tm1*.*Stw*^ mice (n=5 per group). *p<0.05, **p<0.01, one-way ANOVA.

**Figure 7.**
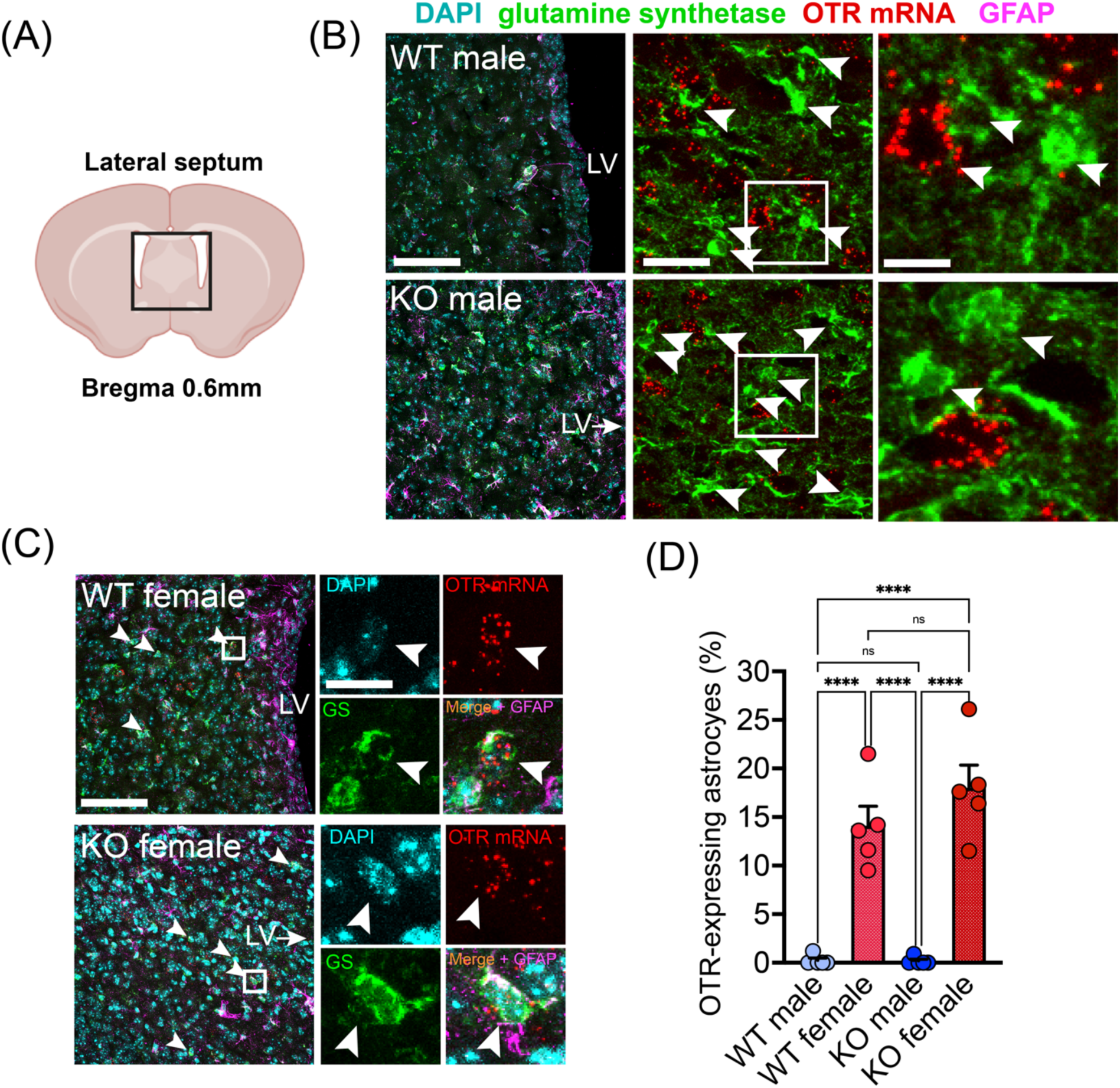
(**A)** Anatomical location of the LS. (**B)** Representative confocal images showing RNAScope hybridization for OTR mRNA and immunostaining against astrocyte markers GS and GFAP in male WT and *Magel2*^*tm1*.*Stw*^ mice (n=5 per group). Arrowheads indicate the mismatch between GS and OTR mRNA signals. LV=lateral ventricle. Scale bar=200µm, 50µm and 10µm. **(C)** Representative confocal images showing RNAScope hybridization for OTR mRNA and immunostaining against astrocyte markers GS and GFAP in female WT and *Magel2*^*tm1*.*Stw*^ mice (n=5 per group). Arrowheads in enlarged insets highlight co-localization between GS and OTR mRNA. Scale bar=200µm and 10µm. **(D)** Quantification highlights the presence of OTR-expressing astrocytes in the LS of female, but not male WT and *Magel2*^*tm1*.*Stw*^ mice. ****p<0.0001, one-way ANOVA.

## 4 Discussion

ASD-like intellectual disability and cognitive deficits are a hallmark of PWS. The hypothalamic neuropeptide OT has been in the focus of PWS research for more than two decades, even since the initial discovery by Swaab and colleagues, which reported a reduction in the number of OT cells within the PVN (10). Given the symptomatic overlap with ASD (56), a plethora of animal and human studies aimed to ameliorate behavioral symptoms via intranasal or systemic application of OT (6, 7, 12-23). However, information about the OTergic system in mouse models of PWS is still scarce and to date, no information about OTR-expressing astrocytes in *Magel2*^*tm1*.*Stw*^ mice exists. To fill this gap, we performed and end-to-end analysis of PVN OT neurons in male and female WT and *Magel2*^*tm1*.*Stw*^ mice using our previously established, semi-automated Imaris pipeline (33, 57). In addition, we assessed OTR mRNA levels in various brain regions, performed detailed analysis of astrocyte numbers and GFAP intensity, as well as assessment of OTR-positive astrocytes in various brain regions.

One intriguing finding is the reduction of OT cells within the caudal part of the PVN, which is in line with early reports from Swaab and colleagues in PWS patients (10), but in stark contrast to previous data from the Muscatelli lab, which actually reported an increase in the number of OT cells (6). While the authors used a higher sample size (n=7-11 mice vs our 4 mice per group), they only analyzed 2-4 sections per brain region and animal according to the supplementary data. In addition, while mice in our current study were 8-12 weeks old, mice in the study from Meziane and colleagues were 3-5 months old. Most importantly, however, Meziane and colleagues used the *Magel2*^*tm1*.*1Mus*^ mouse line, while we used *Magel2*^*tm1*.*Stw*^ line, which could potentially explain the observed differences in the number of OT neurons within the parvocellular subdivision. While the cause for the reduction in OT cells in the caudal part of the PVN remains unclear, it seems possible that *Magel2*^*tm1*.*Stw*^ mice send fewer axons to hindbrain areas such as the dorsal vagal complex or nucleus tractus solitarii, in which OT plays an important role in mediating the satiety effect (11). It is important to note however that while the vast majority of PWS patients develops hyperphagia and obesity, deletion of *Magel2* in *Magel2*^*tm1*.*Stw*^ mice does not reliably cause morbid obesity (8, 9), potentially indicating a contribution of other genes within the PWS locus such as SNORD116 (2). Future studies are clearly needed to investigate the consequence of reduction of OT neurons in the caudal PVN and whether other cell types such as preautonomic vasopressin neurons (58) or corticotropin-releasing hormone neurons (59) are also affected.

When we first discovered OTR-expressing astrocytes in the NAcc, we initially believed that we had made a novel discovery, but after a careful review of existing literature we acknowledge that Gölen and colleagues already briefly described the presence of OTR-bearing astrocytes in the NAcc back in 2013 (see Supplementary Figure S7f in (26). However, the authors did not quantify the number of OTR-positive astrocytes and the fact that lack of *Magel2* protein affects astrocytic OTR levels in this brain region, is a novel finding. Nonetheless, the role of OTR-expressing within these novel brain regions (i.e. LS and NAcc) remains elusive, but it can be speculated that – in analogy to what we described for the CeA – OT might act on astrocytes within these brain regions to facilitate neuromodulatory and behavioral effects, both of which have thus far been exclusively attributed to neuronal OTRs (26, 29). Given the importance of OTergic action in the NAcc during social reward and its intricate interplay with serotonergic circuits and other reward-related brain regions such as the VTA, it is tempting to speculate that activation of OTR-positive astrocytes within the NAcc gates socially-relevant information, which is then transmitted onto the larger population of OTR-expressing neurons. In the LS, OTergic action is needed for maternal behavior, maternal aggression and prevention of social fear in lactating mothers (29-32). Up to this point, the observed effects of OT within this brain region have been solely attributed to OTRs in neurons, but it seems plausible that astrocytic OTRs serve an important, previously unknown function. Whether astrocytic OTR signaling in the NAcc or LS is crucial for proper function of the respective microcircuits and whether OTR-positive astrocytes communicate with neurons via the release of D-serine – as we previously described for the CeA (24) – remains elusive and needs to be addressed in future studies.

In summary, we here present novel findings regarding the oxytocin system and oxytocin receptor-expressing astrocytes in a mouse model of PWS. The consequence for the reduction of OTR-positive astrocytes in the NAcc and the reason for the absence of OTR-positive astrocytes in the LS of male mice needs to be further investigated, but highlights – for the first time – that a genetic condition, namely PWS, could affect OTR-expressing astrocytes in the brain and that sex-specific differences in the number of OTR-expressing astrocytes exist, which could help explain some of the aspects of OT-mediated maternal behavior.

## ACKNOWLEDGEMENTS

The authors thank Dr. Harold Gainer for kindly providing the PS38 antibody. Confocal images presented on Figures 2-7 and S1-3 were acquired at the Nikon Imaging Center, Heidelberg.

## DATA AVAILABILITY

All data generated in this study will be available from the corresponding author upon request.

## SUPPLEMENTAL FIGURES

**Figure S1.**
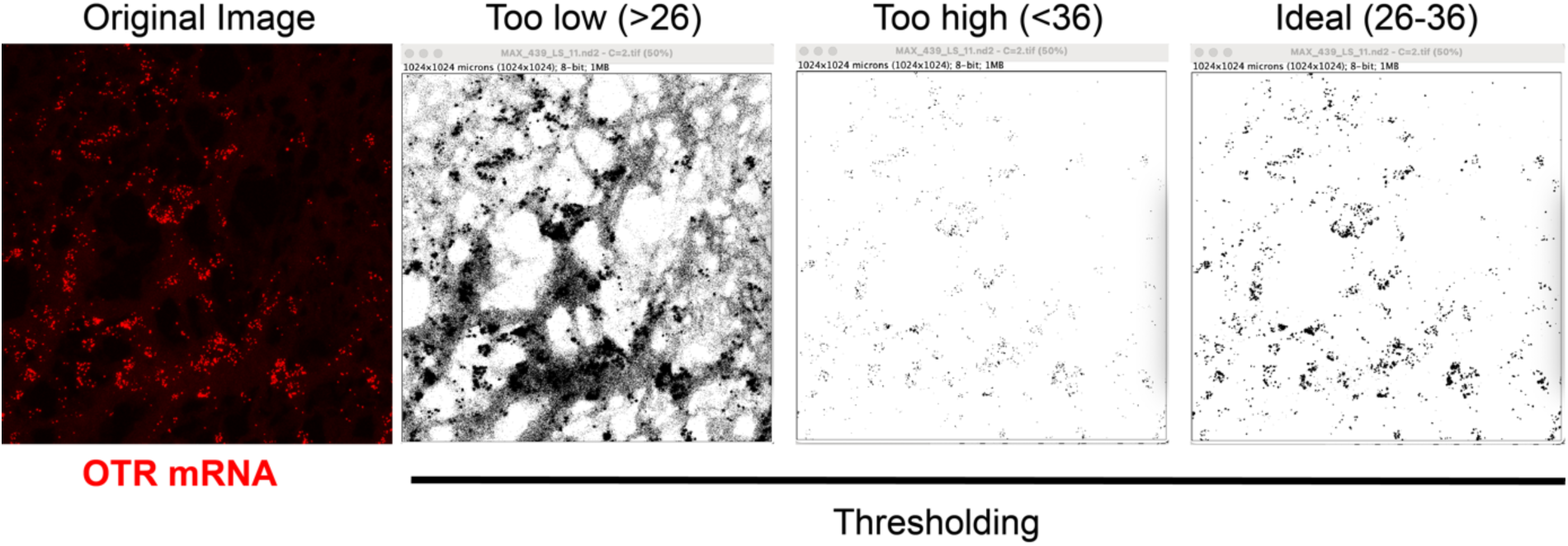
Illustration of the Fiji-based thresholding processing for the OTR mRNA analysis. Raw images were thresholded to achieve near-perfect overlap between actual OTR mRNA signal and binary pixels. All images were processed within the thresholding range of 26-36. Example images show too high, too low and ideal thresholding.

**Figure S2.**
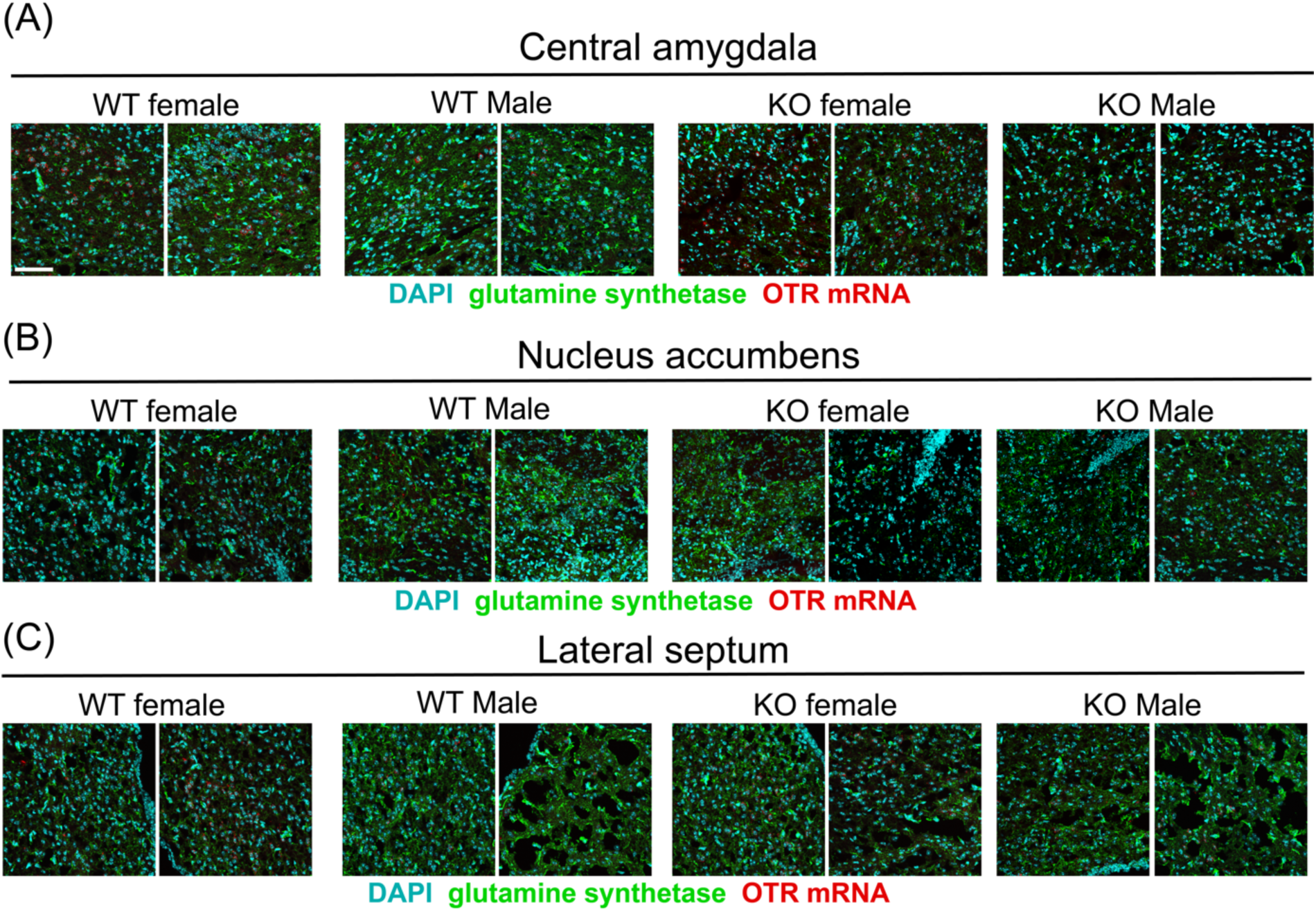
Additional images of RNAScope hybridization showing OTR mRNA (red), GS (green) and DAPI (cyan) in (A) CeA, (B) NAcc and (C) LS in female and male WT and *Magel2*^*tm1*.*Stw*^ mice. Scale bar=100µm.

**Figure S3.**
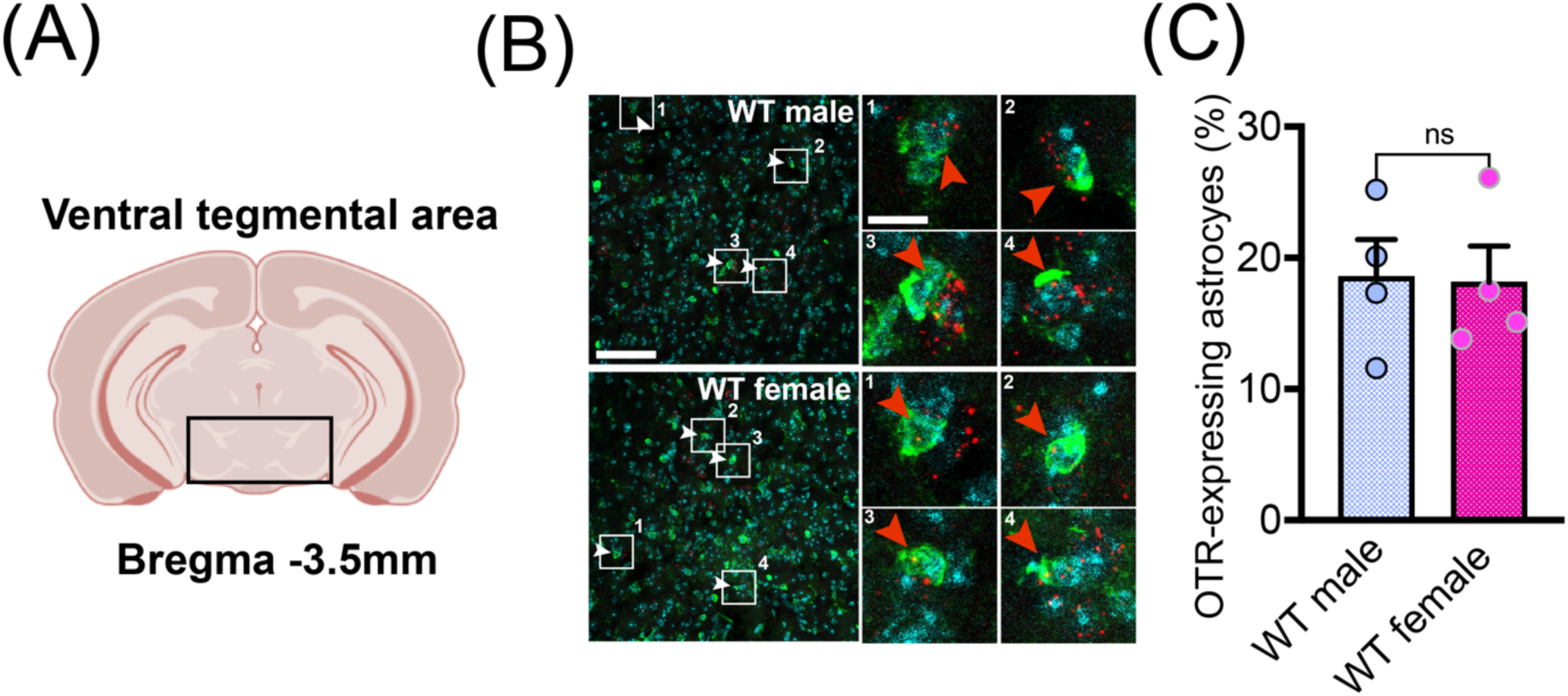
**(A)** Anatomical location of the VTA. **(B)** Confocal images showing OTR-positive astrocytes in the VTA of male and female WT mice. Enlarged insets depict co-localization of GS and OTR mRNA. Scale bar=100µm and 10µm. **(C)** Quantification of OTR-positive astrocytes in male and female WT mice.

